# Cyanobacteria form a procarboxysome-like structure in response to high CO_2_

**DOI:** 10.1101/2024.06.28.601118

**Authors:** Clair A. Huffine, Catherine Fontana, Anton Avramov, Colin Sempeck, Jeffrey C. Cameron

## Abstract

Fixing 25% of CO_2_ globally, cyanobacteria are integral to climate change efforts. The cyanobacterial CO_2_ concentrating mechanism (CCM) features the carboxysome, a bacterial microcompartment which houses their CO_2_ fixing machinery. The proteinaceous shell of the carboxysome restricts diffusion of CO_2_, both inward and outward. While necessary for CCM function in air (0.04% CO_2_), when grown in high CO_2_ levels (3% CO_2_) representative of early earth, the shell would harmfully limit CO_2_ fixation. To understand how carboxysomes change form and function in response to increased CO_2_ conditions, we used a Grx1-roGFP2 redox sensor and single cell timelapse fluorescence microscopy to track subcellular redox states of *Synechococcus* sp. PCC 7002 grown in air or 3% CO_2_. Comparing different levels of compartmentalization, we targeted the cytosol, a shell-less carboxysomal assembly intermediate called the procarboxysome, and the carboxysome. The carboxysome redox state was dynamic and, under 3% CO_2_, procarboxysome-like structures formed and mirrored cytosolic redox states, indicating that a more permeable shell architecture may be favorable when [CO_2_] is high. This work represents a step in understanding how cyanobacteria respond to changing CO_2_ concentrations and the selective forces driving carboxysome evolution.

## Introduction

Photosynthetic bacteria in the phylum cyanobacteria are thought to have reshaped Earth by creating an oxygen-rich atmosphere ∼2.4 Gya; in modern times, they are again poised to significantly alter our planet’s function by serving as a useful carbon dioxide (CO_2_) sink in the face of climate change.^1,2^ Conducting 25% of annual global carbon fixation,^3^ cyanobacteria are crucial to the Earth’s carbon cycle. They accomplish this feat by implementing an efficient CO_2_-concentrating mechanism (CCM) featuring the carboxysome, a proteinaceous bacterial microcompartment housing their carbon-fixing machinery, ribulose-1,5-bisphosphate carboxylase/oxygenase (RuBisCO) and carbonic anhydrase (CA).^4,5^ The carboxysome shell is thought to be selectively permeable to bicarbonate (HCO_3_^-^) while limiting diffusion of molecular oxygen and CO_2_.^6–8^ The CCM also employs numerous membrane-associated HCO_3_^-^ transporters: HCO_3_^-^ is concentrated in the cytosol, diffuses into the carboxysome, is rapidly converted to CO_2_ by CA, and ultimately creates a CO_2_-rich environment around carboxysomal RuBisCO. The permeability of other metabolites and compounds across the carboxysome shell remains largely unexplored. As cyanobacteria originally evolved in a relatively rich CO_2_ environment, they are thought to have developed two convergent lineages of carboxysomes, α/μ, and CCM in response to a simultaneous rise of O_2_ and fall of CO_2_ levels in the atmosphere.^1^ Debate remains in the field on the exact timing and evolutionary pressure of this process.^1,9^ While some caveats do exist, environmental CO_2_ modulation can be used to examine how cyanobacteria may react to increasing CO_2_ levels from climate change as well as explore selective pressures historically experienced by cyanobacteria.^1^

Formation of β-carboxysomes is initiated by aggregation of RuBisCO and CA to the pole of the cell via the scaffold protein, CcmM, into a structure known as the procarboxysome, prior to shell recruitment by CcmN.^10–17^ Since the procarboxysome stage in carboxysome formation represents a transient and short-lived intermediate,^11,18^ little is known on the permeability and functional state of this structure. We are able study the procarboxysome in a perpetual state utilizing shell knock-out mutants. In many cyanobacteria, including *Synechococcus* sp. PCC 7002 (hereafter PCC 7002), the essential trimeric shell protein, CcmO, is encoded at a separate genomic locus distinct from the *ccm*-operon. The *ccm*-operon encodes other necessary carboxysome proteins including the hexameric (CcmK1, CcmK2) and pentameric (CcmL) shell proteins in addition to CcmM and CcmN.^10^ Failure of shell assembly in CcmO knock-out lines (Δ*ccmO*) results in the terminal formation of procarboxysomes (1-2 per cell) without disrupting other core elements of the carboxysome.^10^ *ΔccmO* mutants exhibit a high-CO_2_-requiring (HCR) phenotype and are unable to grow in air (0.04% CO_2_), but can be fully rescued in elevated CO_2_ (3% CO_2_), allowing for study of the procarboxysome directly in high-CO_2_ conditions.^10,18^ By studying the procarboxysome, we can better understand both carboxysome assembly as well as the procarboxysome as a potential evolutionary intermediate.

Reduction-oxidation (redox) regulation is an integral aspect across a number of cellular processes, including function of the CCM and carboxysome (**Fig. 1a**).^15–17,19–22^ Under illumination, cyanobacterial photosynthetic machineries continually generate reactive oxygen species (ROS) through both water splitting and light energy dissipation from pigments. There are three main ROS formed, singlet oxygen (^1^O_2_), hydroxyl radicals (·OH), and hydrogen peroxide (H_2_O_2_). As ROS are both useful as an internal signaling molecules and damaging to the cell, their levels must be carefully regulated. Levels of the longest-lived ROS, H_2_O_2_, are regulated via glutathione (GSH/GSSG), a non-ribosomal peptide-based antioxidant.^23^ GSH is oxidized into GSSG when exposed to H_2_O_2_ and reduced by NADPH with an enzymatic catalyst. In this way, the cytosol is maintained as a reducing environment. Previous work has indicated that the internal redox environment of the carboxysome is an oxidizing environment.^11,20,24^ However, carboxysomal redox state has neither been directly compared to the cytosol nor has a specific redox pool been targeted,^11,20,24^ so much remains to be explored on the redox relationship of the carboxysome to the cytosol under variable conditions. The activity and function of several CCM proteins are known to be redox-regulated, such as one of the HCO_3_^-^ membrane transporters, SbtB/A, as way to modulate carbon uptake,^19,25^ and the scaffold protein, CcmM, as a way to adjust RuBisCO packing during carboxysome formation.^16,17,20–22^ Others have been indicated as redox sensitive, such the shell protein, CcmK4,^21,22^ and both the large and small subunits of RuBisCO, but it is unknown what these redox sensitivities achieve.^21,22^ The purpose and mechanism underlying this distinct redox environment in the carboxysome is still an area of active investigation.

**Fig 1.**
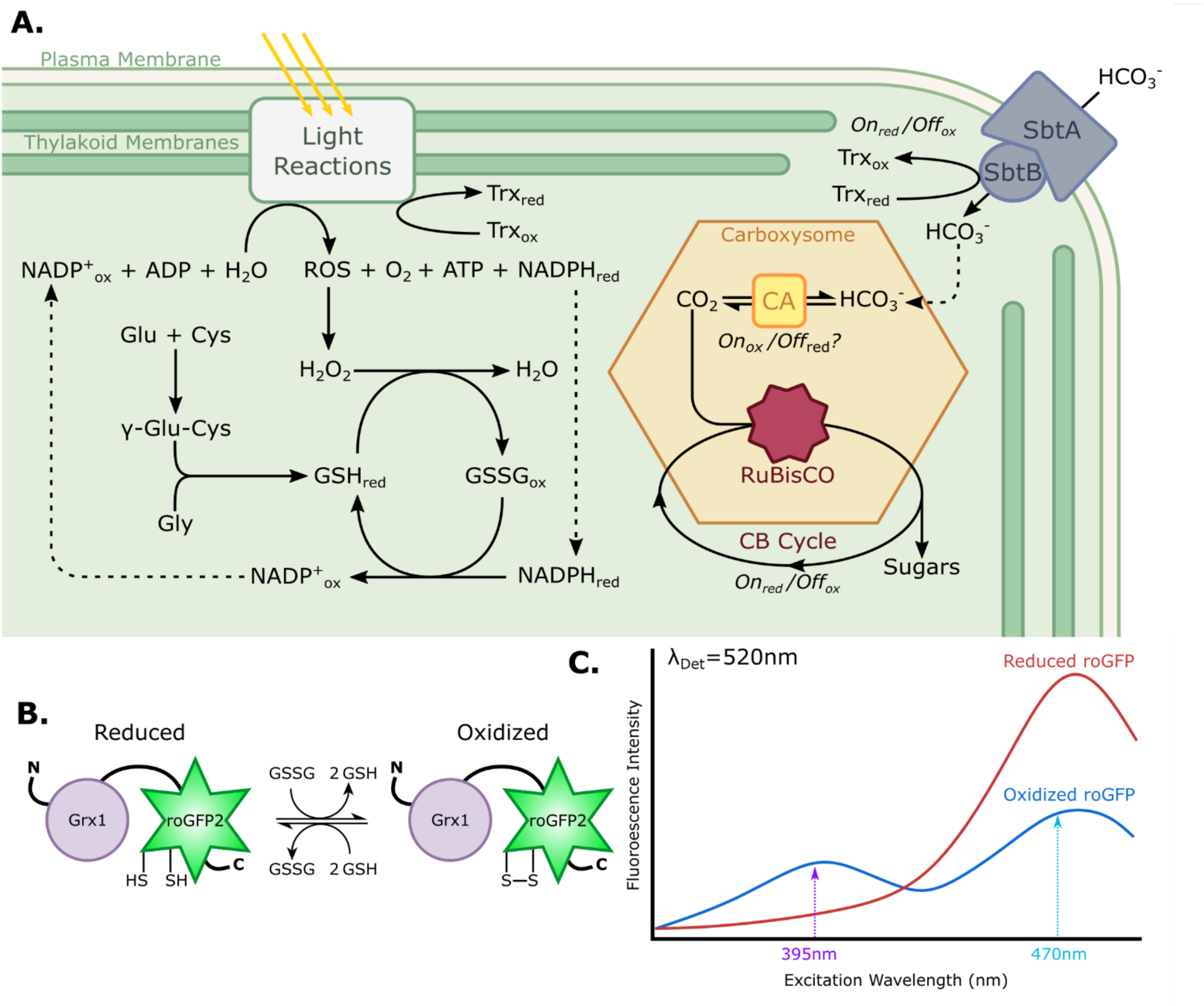
Cyanobacterial CO2-Concentration Mechanism (CCM) and glutathione redox roGFP2 sensor system. (A) Overview of known and theorized redox regulation in cyanobacteria. (B) Grx1-roGFP2 glutaredoxin specifically interacts with glutathione redox pools. (C) Grx1-roGFP2 fluorescence excitation spectrum with capability of ratiometric readout of redox environment.^16^ The bimodal fully oxidized spectrum has a high fluorescence ratio R395/470 compared to the monomodal fully reduced spectrum with a low fluorescence ratio.

While not functional in PCC 7002, in some cyanobacterial strains CcmM has an active γ-CA domain which serves as the carboxysomal CA. This γ-CA has been shown to be redox-regulated.^16,26^ Since a cytosolically located CA would disrupt the HCO_3_^-^ gradient generated by the CCM, CA must be inactivated during carboxysome formation in the cytosol.^27^ The redox regulation of γ-CA in CcmM suggests a clear mechanism by which the γ-CA is able to be inactivated in the reducing cytosol and activated in the oxidized carboxysome via disulfide bond formation. In contrast, the functional CA in PCC 7002 is a β-CA, IcfA (also known as CcaA), for which the regulation is unclear.^28,29^ While pioneer work found this β-CA to be inactivated by reducing agents,^29^ cysteines involved in disulfide bond formation and redox sensing have not been identified.^28^ As the shell-less procarboxysome stage of carboxysome formation contains cytosolically exposed CA, the *ΔccmO* strain provides a unique opportunity to study redox environment in the procarboxysome where CA activity is thought to be inhibited by reduction.

In this work, we track the dynamic redox changes within the cytosol, carboxysomes, and procarboxysomes in PCC 7002 during growth in air (0.04%) and 3% CO_2_ conditions. To accomplish this, we implemented previously characterized redox-sensitive GFPs (roGFP2) fused with glutaredoxin (Grx1) to specifically probe changes in glutathione redox pools (**Fig 1c and d**).^11,24,30,31^ Using single-cell timelapse fluorescence microscopy under precisely controlled environmental conditions,^18,32^ we were able measure relative redox states at subcellular levels in PCC 7002. We discovered previously unknown subcellular redox dynamics in response to elevated CO_2_ conditions, revealing that the carboxysome redox environment is dynamic over time as well as in response to changes in [CO_2_]. This suggests the presence of additional mechanisms for tunable regulation of carboxysome permeability. We also reveal novel procarboxysome-like structures forming in response to elevated CO_2_ conditions. The discovery of procarboxysome-like structures in high CO_2_ could represent a mechanism to overcome the kinetic barrier of the carboxysome shell to CO_2_ and provides insight into the evolutionary pressures leading to the encapsulation of RuBisCO. Furthermore, this study provides a window into how cyanobacteria may adapt to anthropogenetic increases in CO_2_ levels.

## Results

### The Carboxysome is More Oxidized than the Cytosol and Procarboxysome

To probe subcellular cyanobacteria redox environments, we expressed Grx1-roGFP2^30^ either as soluble protein to target the cytosol or as an N-terminal translational fusion with the large subunit of RuBisCO (RbcL) to target the carboxysome/procarboxysome (**Fig 2a**). Live-cell fluorescence imaging showed that the roGFP signal morphology and localization for the carboxysome, cytosol, and procarboxysome was consistent previous literature.^10,18,32^ To assess unintentional disruption of the carboxysome function or shell structure, we tested strains for lack of a high-CO_2_ requiring phenotype.^10,18,33^ Addition of RbcL-Grx1-roGFP2 construct to the *ΔccmO* mutant did not disrupt the existing *ΔccmO* high-CO_2_ requiring growth pattern^10,18^ (**Fig 2b, S1**) while all other strains had WT-like growth in both air and 3% CO_2_ (**Fig 2b, S1**). Overall, the roGFP-expressing strains appeared to be functionally comparable to WT and previously characterized *ΔccmO* mutant^10^ with the growth pattern indicating RbcL-Grx1-roGFP2 addition did not disrupt normal carboxysome function or localization.

**Fig 2.**
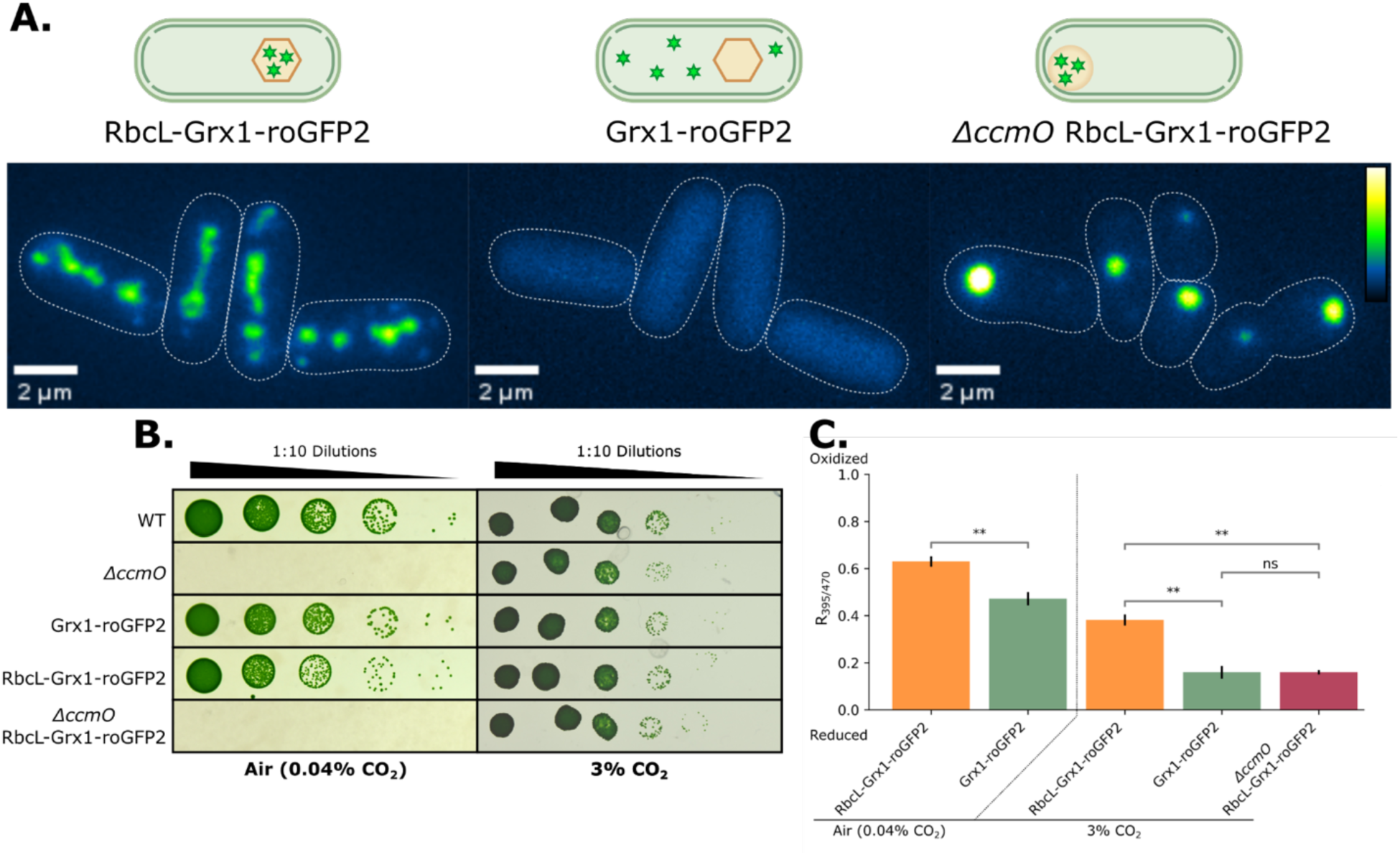
Grx1-roGFP2 Strains Characterization. (A) Fluorescence microscopy images of GFP intensity and localization in exponentially growing PCC 7002 strains growing in air (RbcL-Grx1-roGFP2 and Grx1-roGFP2) or growing in 3% CO2 (*ΔccmO* RbcL-Grx1-roGFP2). Indicating the localization pattern of Grx1-roGFP2 to each of the subcellular regions of interest (carboxysome, cytosol, and procarboxysome, respectively). Color bar indicates GFP intensity. (B) Growth of strains in air (0.04% CO2) and 3% CO2 on 1% and 0.5% agar spot plates, respectively.^33^ RoGFP strains showed either WT or *ΔccmO* like growth. Ten-fold serial dilutions were plated and imaged after 72 hours. Images are representative of three biological replicates. (C) Spectrofluorometer measurements of strains grown in bulk liquid in air and 3% CO2. The carboxysome (RbcL-Grx1-roGFP) is more oxidized than the cytosol (Grx1-roGFP) and procarboxysome (*ΔccmO* RbcL-Grx1-roGFP2) and both the cytosol and carboxysome become more reduced under 3% CO2 than in air. No data is included for the procarboxysome redox state under air as *ΔccmO* RbcL-Grx1-roGFP2 cannot grow in air. Cultures were grown in respective CO2 conditions for 24 hours prior to measurement. Normalized prior for chlorophyll content, cells were excited from 350 nm to 480 nm with emission intensity collected at 510 ± 5 nm and WT emission signal was subtracted from each corresponding growth condition to account for background emission. The results are representative of three biological replicates, n = 3, bars represent one standard deviation, and data was analyzed by Student’s t-test. *, p < 0.05; **, p < 0.01.

To confirm sensitivity of the roGFP probe, oxidizing (H_2_O_2_) and reducing agents (DTT) were added to bulk cultures and the fluorescence emission spectra of roGFP was measured, generating ratiometric readouts of the redox environment in each strain (**S2a**). The cytosol showed consistent redox responses whereas the carboxysome was unresponsive to H_2_O_2_ addition, perhaps indicating differences in shell permeability of the oxidizing and reducing agents. No significant changes were measured in the procarboxysome with either redox agent, likely due to the preexisting reduced state from growing in 3% CO_2_ limiting DTT impact and lower sensitivity to H_2_O_2_ as seen in the carboxysome (**S2b**). Because the roGFP probe relies on a ratiometric measurement, the measurement of redox state is independent of roGFP concentration, subsequently allowing for comparison of strains with differing GFP signal intensity.^30^ This feature of the roGFP system is supported by comparable redox readouts of the diffuse cytosol signal and the cytosolically exposed procarboxysome with high intensity puncta (**Fig 2a and c**). These results indicate that the roGFP system is functional at a bulk culture level and can be used to probe the redox poise of different subcellular regions.

To observe if subcellular redox states respond to [CO_2_] changes, the redox state was measured in bulk cultures grown either in air or 3% CO_2_ (**Fig 2c, S2**). Carboxysomes were more oxidized than the cytosol in both air and 3% CO_2_ (**Fig 1c**), however, unexpectedly, both the cytosol and carboxysomes became more reduced in 3% CO_2_ conditions compared to their air-grown counterparts. Although previous work has probed the carboxysome environment independent of the cytosol,^11,24^ this is the first time, to our knowledge, that the redox environment of these two subcellular regions have been directly compared. The procarboxysome redox state was not significantly different than the cytosol in 3% CO_2_. This supports that the procarboxysome, with its non-existent shell,^10^ is exposed to the cytosolic environment with a similar reduced redox environment. Furthermore, CO_2_ concentration impacts redox environment in subcellular regions of PCC 7002.

### The Carboxysome Redox State Dynamically Responds to [CO_2_]

The bulk culture results revealed that the redox state in cyanobacteria responded to [CO_2_]. To investigate this response to 3% CO_2_ at a finer scale, we used timelapse microscopy to capture the redox dynamics of subcellular regions of PCC 7002. This approach allowed for simultaneous comparison of redox dynamics across a population as well as individual cell responses. In agreement with the bulk data, the carboxysome was consistently more oxidized than the cytosol and procarboxysome (**Fig 3a-e**). When the growing conditions were changed from air to 3% CO_2_, there was an approximately 8-hour shift of the carboxysome to more reduced redox environment (**Fig 3f-j**). However, when reversed from 3% CO_2_ back to air, the average redox shift of the carboxysome to a steady-state more oxidizing environment exhibited hysteresis and did not return to the same pre-high CO_2_ redox state (**S3**). These observations suggest that the redox state in the carboxysome is dynamic, but, at the average population level, did not explain what might drive these shifts.

**Fig 3.**
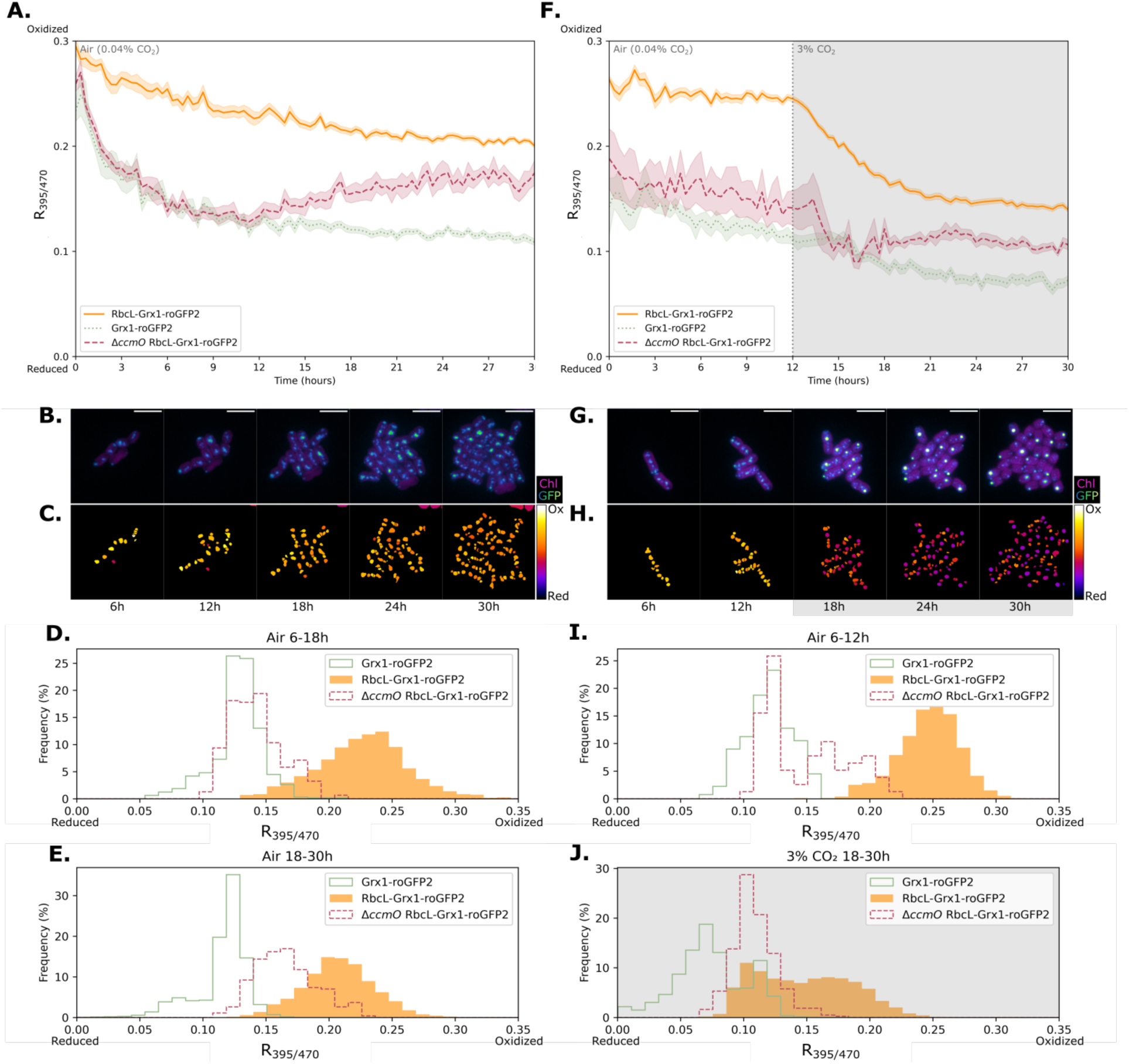
Redox state dynamics under modulated [CO2]. Aggregated redox state from timelapse fluorescence microscopy of the carboxysome (RbcL-Grx1-roGFP2), procarboxysome (*ΔccmO* RbcL-Grx1-roGFP2), and cytosol (Grx1-roGFP2) over 30 hours of growth in (A-E) air or (F-J) 3% CO2 conditions from hour 12 to 30. (B and G) GFP and chlorophyll fluorescence and (C and H) ratiometric images of representative carboxysomes over 30 hours of growth. Redox color bar spans from *R*_395/470_ to 0.3. Histograms represent frequency of redox state of each subcellular region when growing in air (D, E, I) or 3% CO2 (J). Wildtype background fluorescence was subtracted from excitation intensity values at 395 nm and 470 nm (emission at 520 nm). Error bars represent standard error with n changing over the course of the experiment for each strain as a result of cell division, (A) n ≥ 6, and (F) n ≥ 4 to n ≥ 18. Representative data from three biological replicates.

### Procarboxysome-Like Structures Form in 3% CO_2_

Further analysis of the population abundances of subcellular redox states revealed that in 3% CO_2_ the carboxysomes in RbcL-Grx1-roGFP2 have a bimodal distribution (**Fig 3j, S3**) which disappears when returned to air (**S3**). In addition, we also noticed changes in the morphology of the fluorescent puncta labeling carboxysomes in RbcL-Grx1-roGFP2. When grown in 3% CO_2_, there was formation of large, high intensity signal puncta with redox states comparable to the procarboxysome puncta in *ΔccmO* RbcL-Grx1-roGFP2 (**Fig 3**, **Fig 4a**) which we have termed procarboxysome-like structures. Once the cells were returned to air, the reduced state of these procarboxysome-like structures persisted for <6 hours before assumably being processed into carboxysomes (**S3**). Given that the procarboxysome and procarboxysome-like structures share a similar redox state with the cytosol (**Fig 3j, S3**) and that the procarboxysome does not have a shell,^10^ we therefore concluded that the procarboxysome-like structures must also have either a limited or completely absent shell.

**Fig 4.**
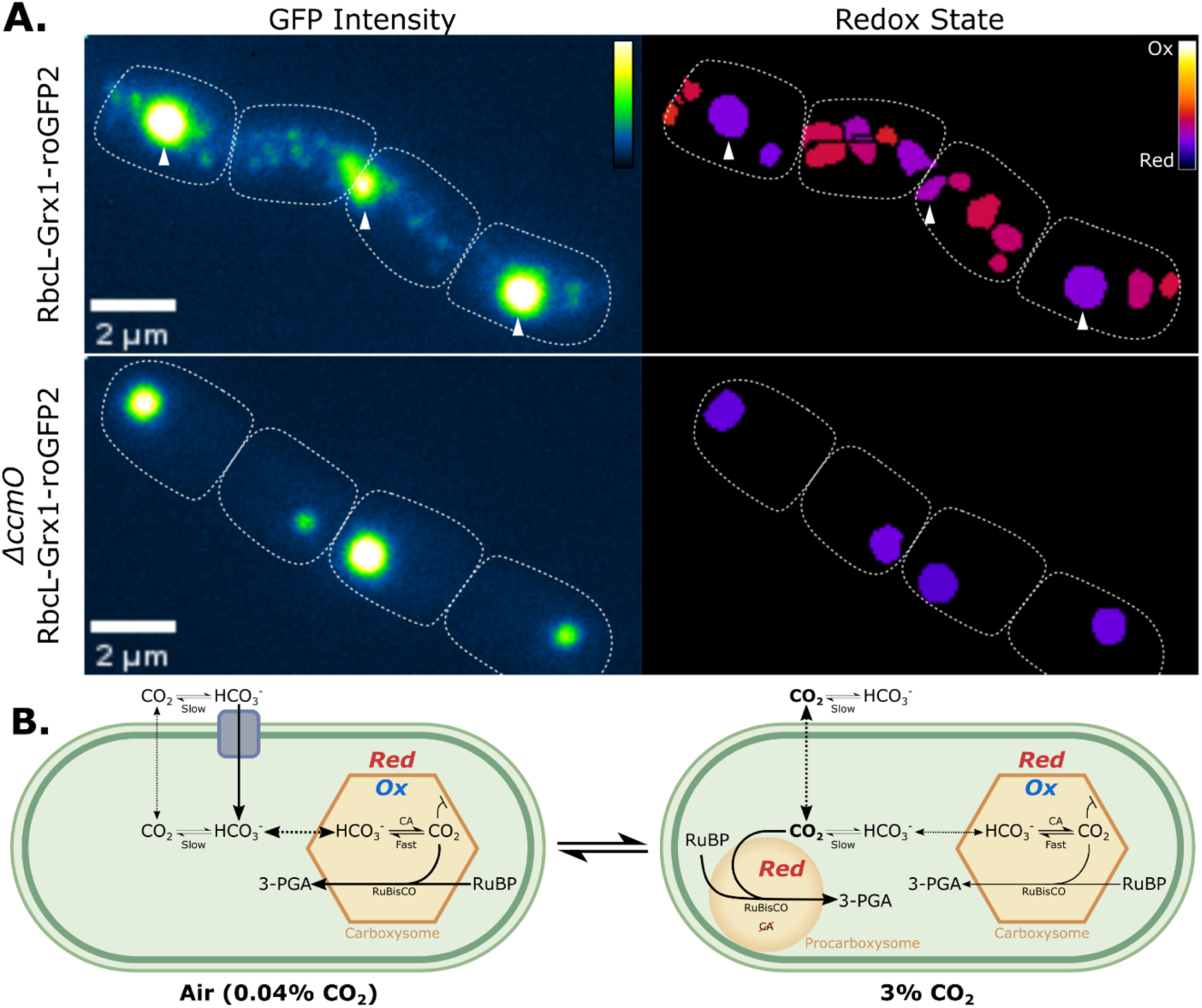
Procarboxysome-like Structures form in 3% CO2. (A) GFP fluorescence microscopy showing that under 3% CO2 there are procarboxysome-like puncta present in the carboxysome-labeled strain (RbcL-Grx1-roGFP). These procarboxysome-like structures have a similar size, fluorescent intensity, and redox state as procarboxysomes as seen in the procarboxysome-labeled strain (*ΔccmO* RbcL-Grx1-roGFP2). Color bars indicate GFP intensity and redox ratio (0-0.5) respectively (B) Proposed dynamic cyanobacterial CCM model under low and high CO2 conditions. Under reducing environments carbonic anhydrase (CA) activity is thought to be inhibited,^29^ and bicarbonate (HCO3^-^) transport downregulated as the pH shift in 3% CO2 reduces [HCO3^-^]. Given the theorized impermeability of the carboxysome shell to CO2 it would be beneficial to have RuBisCO unencapsulated under high CO2 conditions.

## Discussion

### Dynamics of Carboxysome Redox State

This work represents the first exploration on the impact of [CO_2_] on the subcellular cyanobacteria redox environment. The dynamic redox state, exemplified by a shift of carboxysomes to a more reduced environment over time (**Fig 3a-e**), brings up the unanswered question of what drives and maintains carboxysome oxidation state in the first place. While redox regulation has repeatedly been implicated in controlling γ-CA activity^16,26^ and carboxysome aggregation via CcmM structure and binding affinity,^16,17,20^ to our knowledge, there has not yet been an identified component of the carboxysome system capable of actively oxidizing the internal carboxysome environment during carboxysome formation. We speculate that what drives carboxysome oxidation could be an unidentified oxidizing protein localized to the carboxysome or a gradually oxidizing trapped glutathione pool resulting from shell impermeability to reducing agents or protein catalysts.

None of the known carboxysome components have been reported to possess enzymatic oxidizing capability. It is possible that the presence of the roGFP sensor alters the redox environment in which it is located and artificially creates an oxidized carboxysomal environment but, given the evidence for redox regulation of CAs and CcmM, there is biologically based functional support for the carboxysome being an oxidized environment. Other redox probes could be implemented but given the unknown permeability of the carboxysome shell to chemical probes, this nanometer-scale subcellular region^34^ remains challenging to study.

There was a consistent shift in redox environment the first 2-3 hours, likely a consequence of the cells adjusting to the environmental conditions of the microscope and therefore was disregarded in the CO_2_ modulation data.^33^ However, this initial shift still has intriguing implications. Since the state of the carboxysome appears to start out oxidized then trend towards a more reduced steady state over time (**Fig 3a and f, S3**), it is unlikely that the glutathione pool trapped during carboxysome formation oxidizes from inability to be reduced by cytosolically located NADPH alone. Additionally, the dynamic redox environment of the carboxysome suggests potential for an adjustable permeability of the carboxysome to redox agents, such as DTT (**S2a**). Further work is needed to explore carboxysome redox dynamics, shell permeability to redox agents, and processes driving carboxysome oxidation.

### Formation of Procarboxysome-like Structures

Discovery of procarboxysome-like structures forming when cells are in CO_2_ conditions provides both insights into carboxysome evolution and function and opens new questions. Unique to the work presented here, we leveraged the terminal procarboxysomes in *ΔccmO* mutants to directly compare similarities in morphology and redox state of the procarboxysome-like structures forming high [CO_2_] in non-knockout cells. We hypothesize that since the carboxysome shell is thought to be a CO_2_ barrier,^6,7^ as [CO_2_] increases, it becomes detrimental to have exclusively encapsulated RuBisCO (**Fig 4b**). This is supported by previous findings in which the photosynthetic capacity of PSII, which is sensitive to [CO_2_], is increased in strains without carboxysomes, but only in elevated CO_2_.^10^ The exact mechanism of redox (in)activation of β-CA needs to be more deeply explored in both PCC 7002 and other β-carboxysome cyanobacteria, but CA may be inactivated in response to being in a more reduced state in high [CO_2_] conditions (**Fig 4b**).^29^ As a result, procarboxysome-like structure formation would be preferential as RuBisCO can remain exposed to the higher CO_2_ without needing the CCM and CA activity.

By tracking cytosol redox state in cyanobacteria, we were able to further support that procarboxysomes and the procarboxysome-like structures share a reduced state like the cytosol. Given that procarboxysome-like structures are exposed to the cytosol and do not rapidly proceed to forming carboxysomes once returned to air, there is some delayed process involved in carboxysome formation. These procarboxysome-like structures could have enough excess RuBisCO aggregated that shell proteins, with decreased expression in high [CO_2_] conditions,^35–38^ cannot immediately encapsulate the RuBisCO into carboxysomes when returned to low [CO_2_].^10,18^ Alternatively, if this mechanism is not driven solely by changes in shell protein production, perhaps there is a RuBisCO structural shift once it has been activated via carbamylation in high CO_2_^39^ or some other persisting signaling factor that delays carboxysome formation.

High CO_2_-specific formation of procarboxysome-like structures could point to the procarboxysome as an evolutionary intermediate during changing CO_2_ conditions.^1^ Studies on *Synechococcus* sp. PCC 7942^40^ and *Synechocystis* sp. PCC 6803^34,37^ grown in increased [CO_2_] found there were less carboxysomes, but increased CO_2_ fixation productivity supporting down regulation of carboxysome formation under these conditions. While unable to be identified at the time as procarboxysomes or procarboxysome-like structures, work from pioneer studies observed that a subset of carboxysomes in *Synechococcus* UTEX 625 grown in 5% CO_2_ were larger and irregularly shaped,^41^ indicating that perhaps formation of procarboxysome-like structures in high [CO_2_] is not a strain specific phenomena.

This work paves the way for a more detailed understanding of carboxysome formation, function under altered environmental conditions, and redox regulation of carbon fixation. By better understanding these processes, we can more effectively implement the CO_2_ capturing ability of cyanobacteria for climate change mitigation and biofuel production as well as guide research on the pressures driving carboxysome evolution.

## Methods

### a. Strain cultivation

PCC 7002 strains were cultivated in AL-41 L4 Environmental Chambers (Percival Scientific, Perry, IA) at 37°C under constant illumination (∼150 µmol photons m^−2^ s^−1^) by cool white, fluorescent lamps, under either ambient (air, 0.04%) or elevated (3%) CO2 conditions. Cultures were grown in 25 ml of A+ media in orbital shaking baffled flasks (125 ml) contained with foam stoppers (Jaece Identi-Plug), or on pH 8.2 A+ media solidified with Bacto Agar (1%; w/v). Antibiotics were added for routine growth of strains (kanamycin, 100 µg/ml; gentamycin, 30 µg/ml).

### b. Plasmid and strain construction

All plasmids and strains used in this work are described in Table S1 and Table S2. Plasmids were created through Gibson assembly of plasmid backbones (pUC19) and PCR-amplified inserts, generated using Phusion polymerase (Thermo Fisher Scientific) and primers described in Table S3. Cyanobacterial strains were generated by transforming cells in exponential/early linear growth phase with 0.5 ng/ml of plasmid DNA, containing the insert of interest flanked by 600–base pair homology arms for recombination into a specified genomic locus. After incubation at 30°C in constant illumination (50 to 150 µmol photons m^−2^ s^−1^) for 24 hours, transformed cells were selected for with appropriate antibiotic on plates in ambient CO2, for non-high-CO2 requiring strains, and 3% CO2 for high-CO2 requiring strains, respectively. From plates, individual colonies were patched onto new plates and tested for segregation. Confirmation of segregation was confirmed by PCR, using primers specific for *glpK*. Presence of the insert-specific PCR product and absence of the WT-specific PCR product was used as an indicator of full segregation.

### c. Spot plating

The growth of PCC 7002 was measured on agar plates as described. Plates at 0.5 and 1% agar were spotted with strains in triplicate. Liquid cultures of each PCC 7002 strain were diluted to 0.05 OD730 and five 1:10 serial dilutions were performed. Five µL of the serial dilutions was used for each spot and allowed to dry (30 min) prior to incubation. Images were taken 3 days after spotting the plates with a backlight on a Kaiser eVision light plate and imaged with a Nikon D7200 digital single-lens reflex camera.

### d. Liquid Growth Curves

The growth of PCC 7002 was measured in liquid cultures as described. The precultures were started from PCC 7002 cells scraped from plates and grown in the same conditions as the growth curve cultures, either ambient (air, 0.04%) or elevated (3%) CO2 conditions. 50mL A+ cultures were inoculated in triplicate with 1 mL of PCC 7002 pre-culture diluted to 0.14 OD730 and grown in the standard conditions described in Strain Cultivation. During the growth curve, time points were taken every 24 hours for 72 hours. At each time point, 200 µL was removed from each culture and the OD730 was measured in a 96-well plate on a Tecan Spark multimode microplate reader.

### e. Spectrofluorometer

#### i. Chlorophyll quantification

50mL cultures were inoculated from pre-cultures grown in liquid cultures. Liquid cultures were grown to OD730 0.3-1.0 in either ambient (air, 0.04%) or elevated (3%) CO2 conditions. Chlorophyll was methanol extracted from 1 mL of culture diluted to 0.3 OD730 as described in Porra et al..^42^ Absorbance at 665 nm was measured and the chlorophyll content was calculated with equation 1.

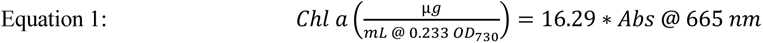

#### ii. Fluorescent Spectra Measurement

Once each culture’s chlorophyll had been quantified, each original culture was diluted to a chlorophyll concentration of 3 µg/mL in A+ media. The normalized chlorophyll cultures were loaded into a FireflySci 1FLPS Disposable Cuvette. Fluorescence was measured using a Fluorolog-3 spectrofluorometer (Horiba Jobin Yvon). Grx1-roGFP2 was excited from 350- to 480 nm with a 5 nm slit and a step size of 1 nm and the fluorescence emission spectra was gathered with an emission wavelength of 510 nm with a 5 nm slit. For sensitivity tests, 30 mM H2O2 or 100 µM DTT was added to the cuvettes and allowed to incubate for 30 s prior to measurement.

#### iii. Ratiometric Data Processing

In replicates of three or four, WT emission was averaged at excitation at 395- and 470 nm respectively (b395 and b470). This value was then subtracted from each Grx1-roGFP2 strain’s emission value from 395- and 470 nm excitation respectively (I395 and I470) before dividing the emission from 395 nm excitation by the emission from 470 nm excitation (equation 2) and averaging across samples.

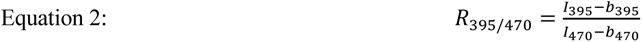

### f. Quantitative microscopy

Fluorescence images were taken using a customized Nikon TiE inverted wide-field microscope with a near-infrared–based Perfect Focus System.^18,32^ Temperature and CO2 concentrations were controlled with a Lexan environmental chamber outfitted with a ProCO2 P120 Carbon Dioxide Single Chamber Controller (BioSpherix, Parish, NY), and growth light was controlled via a transilluminating red light emitting diode (LED) light source (Lida Light Engine, Lumencor, Beaverton, OR). A highspeed light source with custom filter sets was used for imaging (Spectra X Light Engine, Lumencor, Beaverton, OR), along with a hardware-triggered and synchronized shutter for control of imaging and growth light. NIS Elements AR software (version 5.11.00 64-bits) with Jobs acquisition upgrade was used to control the microscope. Image acquisition was performed using an ORCA-Flash4.0 V2+ Digital sCMOS camera (Hamamatsu) with a Nikon CF160 Plan Apochromat Lambda 100× oil immersion objective (1.45 numerical aperture).

For long-term time-lapse microscopy, cells in exponential or early linear phase were diluted to 0.14 OD730, all strains were mixed in equal proportions, and 1 µL was spotted onto a 1% agarose A+ pad. Cells were dried onto the pad (20 min), inverted onto a 35-mm glass bottom imaging dish (ibidi), which was then wrapped in parafilm to keep the pad from drying out, and preincubated at 37°C for 1 hour in the dark. No antibiotics were included on the agarose pad. Images were taken every 20 min using a 395-, 470-, 555- and 640 nm LED light source (Spectra X) and emission wavelengths were collected using standard GFP (395- and 470 nm excitation, 520 nm emission), RFP (595 nm emission), and Cy5 (705 nm) filters (Nikon). Cells were constantly illuminated with red light except during fluorescent imaging.

### g. Image processing and analysis

Cell segmentation was performed using MATLAB version R2020b as outlined previously.^32^ To segment (identify) individual cells, we also captured images in bright field, with the red growth light as an illumination source. Cells were then identified by applying an intensity threshold and watershed algorithm to create a cell mask. Manual mask correction was then performed to correct mistakes before data analysis. Cells that died or overlapped were removed from the mask and subsequent data analysis. Carboxysome and procarboxysome puncta were further segmented based on their GFP signal. Note that these mask images were only used for cell segmentation—reported data were measured from the original images.

Each cell’s strain was visually identified. Puncta smaller than 62 pixels in the *ΔccmO* mutant were excluded from analysis to limit misidentified puncta from background noise. Averaged intensity of WT was used for background subtraction for 395- and 470 nm excitation channels from the averaged intensity of each cell or puncta. To account for low signal in the 470 nm excitation channel, any cell or puncta that was below zero after background subtraction was brought to zero for subsequent calculations. Redox states were calculated across all strains using Equation 2. This ratio was overlayed on the respective cell or puncta mask to generate ratiometric images for ease of visualization.

### h. Statistics

For the statistical comparison of *R*_395/470_ for bulk culture redox state, unpaired two-tailed Student’s t-tests were used. P values are indicated by asterisks; *p < 0.05, **p < 0.001, ***p < 0.0001.

## Author Contributions

J.C.C. and C.A.H. conceived the project. C.A.H., C.F., A.A., and C.S. performed experiments and analyzed data. J.C.C. and C.A.H. wrote the manuscript with input from all the authors.

## Supplementary Data

**Supplementary Figure 1.**
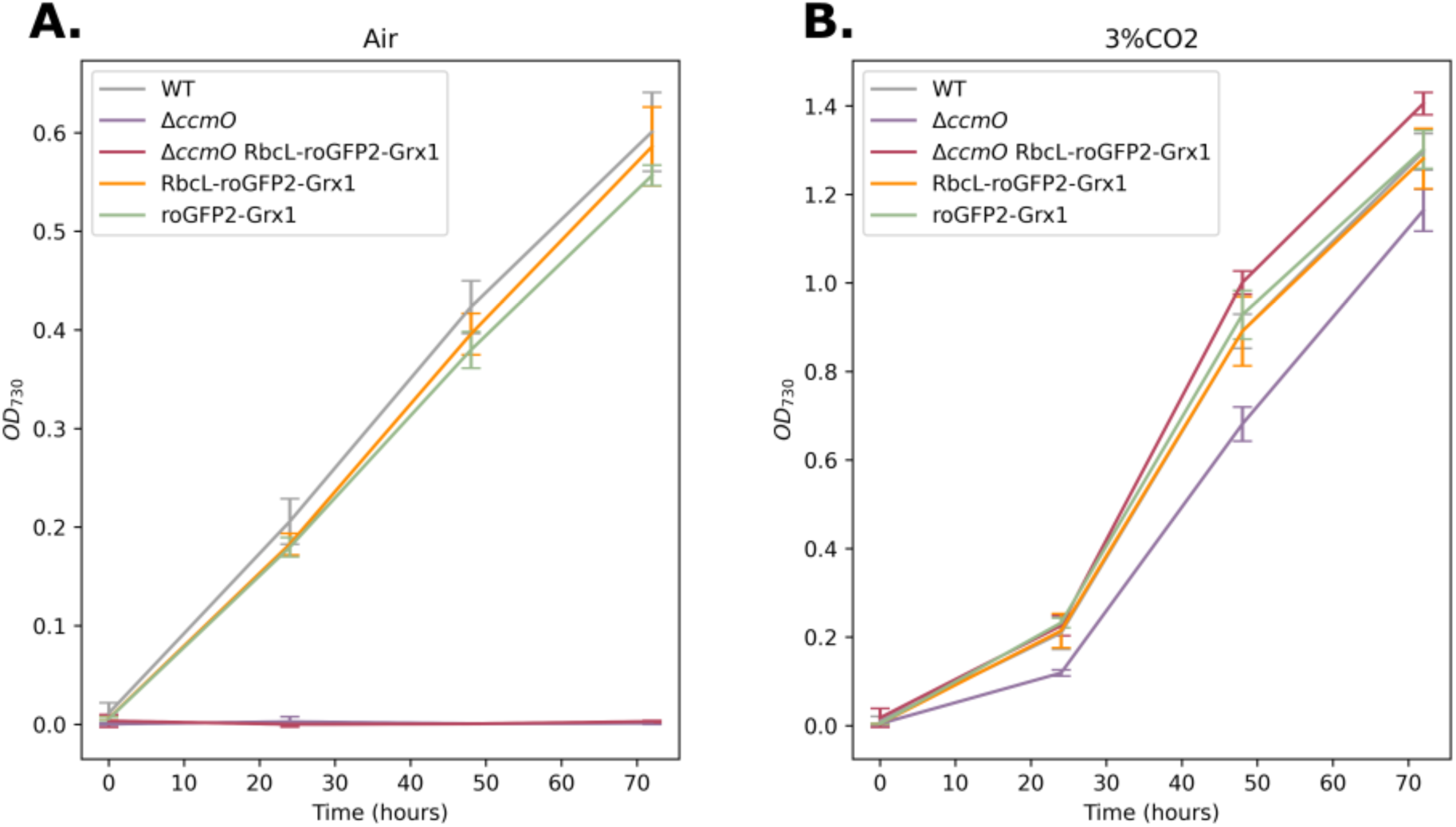
Liquid Growth Curves. (A) Liquid growth curves in air or (B) 3% CO2 showing WT-like growth for all strains in air and 3% CO2 besides *ΔccmO* RbcL-Grx1-roGFP2. *ΔccmO* RbcL-Grx1-roGFP2 was unable to grow in air but growth was rescued in 3% CO2. n = 3, error bars represent one standard deviation.

**Supplementary Figure 2.**
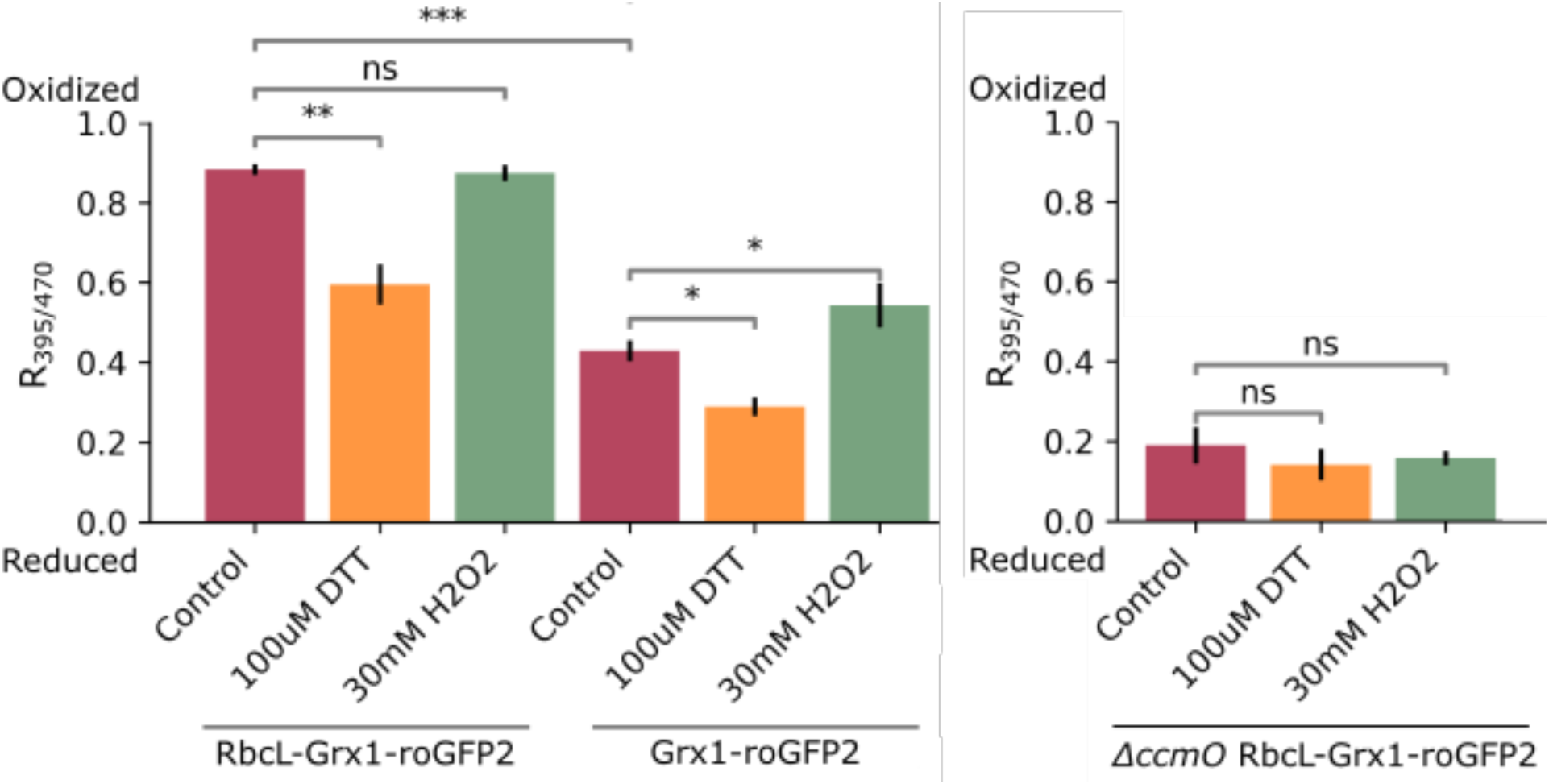
Bulk Redox Sensitivity. Spectrofluorometer measurements of strains grown in bulk liquid with 100 µM DTT and 30 mM H2O2 added 25 s prior to measurement. (A) Grown in air, the carboxysome (RbcL-Grx1-roGFP) and the cytosol (Grx1-roGFP) are reduced when exposed to DTT whereas only the cytosol is oxidized with exposure to H2O2. (B) Grown in 3% CO2, the procarboxysome (*ΔccmO* RbcL-Grx1-roGFP2) does not change redox state under either condition. Cultures were grown in respective CO2 conditions for 24 hours prior to measurement. Normalized prior for chlorophyll content, cells were excited from 350 nm to 480 nm with emission intensity collected at 510 ± 5 nm and WT emission signal was subtracted from each corresponding growth condition to account for background emission. The results are representative of three biological replicates, n = 3, bars represent one standard deviation, and data was analyzed by Student’s t-test. *, p < 0.05; **, p < 0.01.

**Supplementary Figure 3.**
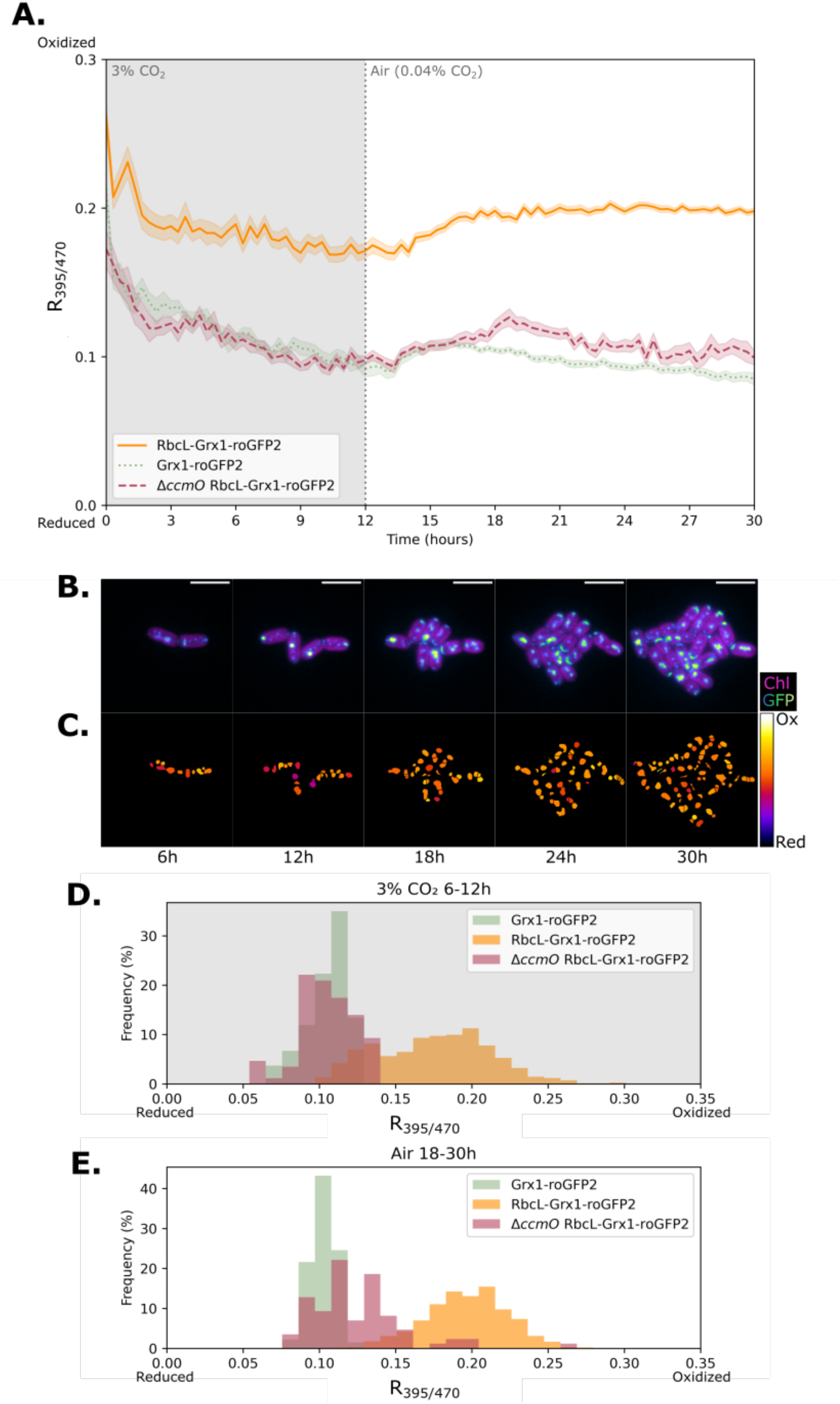
High to Low [CO_2_] Redox Timelapse. (A) Aggregated redox state from timelapse fluorescence microscopy of the carboxysome (RbcL-Grx1-roGFP2), procarboxysome (*ΔccmO* RbcL-Grx1-roGFP2), and cytosol (Grx1-roGFP2) over 30 hours of growth in 3% CO2 conditions from hour 0 to 12. (B) GFP and Chlorophyll fluorescence and (C) ratiometric images of representative carboxysomes over 30 hours of growth. Redox color bar spans from 0 to 0.3. Histograms represent frequency of redox state of each subcellular region when growing in (D) 3% CO2 or (E) air. Wildtype background fluorescence was subtracted from excitation intensity values at 395 nm and 470 nm (emission at 520 nm). Error bars represent standard error with n changing over the course of the experiment for each strain as a result of cell division, n ≥ 5 to n ≥ 24. Representative data from three biological replicates.

**Supplementary Table 1.** Strains used in this study.

| Name | Number | Description | Resistance | Reference |
| --- | --- | --- | --- | --- |
| WT | - | Wild-type *Synechococcus* sp. PCC 7002 | None | - |
| Grx1-roGFP2 | scJC0501 | WT cells transformed with sJC0694 | Gm<sup>R</sup> | This work |
| RbcL-Grx1-roGFP2 | scJC0479 | WT cells transformed with sJC0683 | Km<sup>R</sup> | This work |
| *ΔccmO* | scJC0079 | WT cells transformed with sJC0132 | Km<sup>R</sup> | This work |
| *ΔccmO* RbcL-Grx1-roGFP2 | scJC0503 | *ΔccmO* cells transformed with sJC0691 | Km<sup>R</sup>, Gm<sup>R</sup> | This work |

**Supplementary Table 2.** Plasmids used in this study.

| Name | Number | Description | Genomic Site | Reference |
| --- | --- | --- | --- | --- |
| Grx1-roGFP2-His | sJC0658 | Used as Grx1-roGFP2 PCR template to make sJC0694 and sJC0683 | - | Gutscher 2008<sup>30</sup> |
| Grx1-roGFP2-His_Gm<sup>R</sup> | sJC0694 | PK2_Grx1-roGFP2 downstream of Km<sup>R</sup> cassette | *glpK* | This work |
| RbcL-Grx1-roGFP2-His_Km<sup>R</sup> | sJC0683 | PK2_RbcL-Grx1-roGFP2 downstream of Km<sup>R</sup> cassette | *glpK* | This work |
| RbcL-Grx1-roGFP2-His_Gm<sup>R</sup> | sJC0691 | PK2_RbcL-Grx1-roGFP2 downstream of Gm<sup>R</sup> cassette | *glpK* | This work |
| *ΔccmO*_Km<sup>R</sup> | sJC0132 | Km<sup>R</sup> cassette flanked by *ccmO* operon upstream and downstream homology arms | *ccmO* | This work |
| RbcL-GFP-V5_Km<sup>R</sup> | sJC0624 | Used as a PCR template to make cJC0683 | *glpK* | This work |
| RbcL-GFP-APEX2_Gm<sup>R</sup> | sJC0490 | Used as a PCR template to make cJC0691 and cJC0694 | *glpK* | This work |

**Supplementary Table 3.**
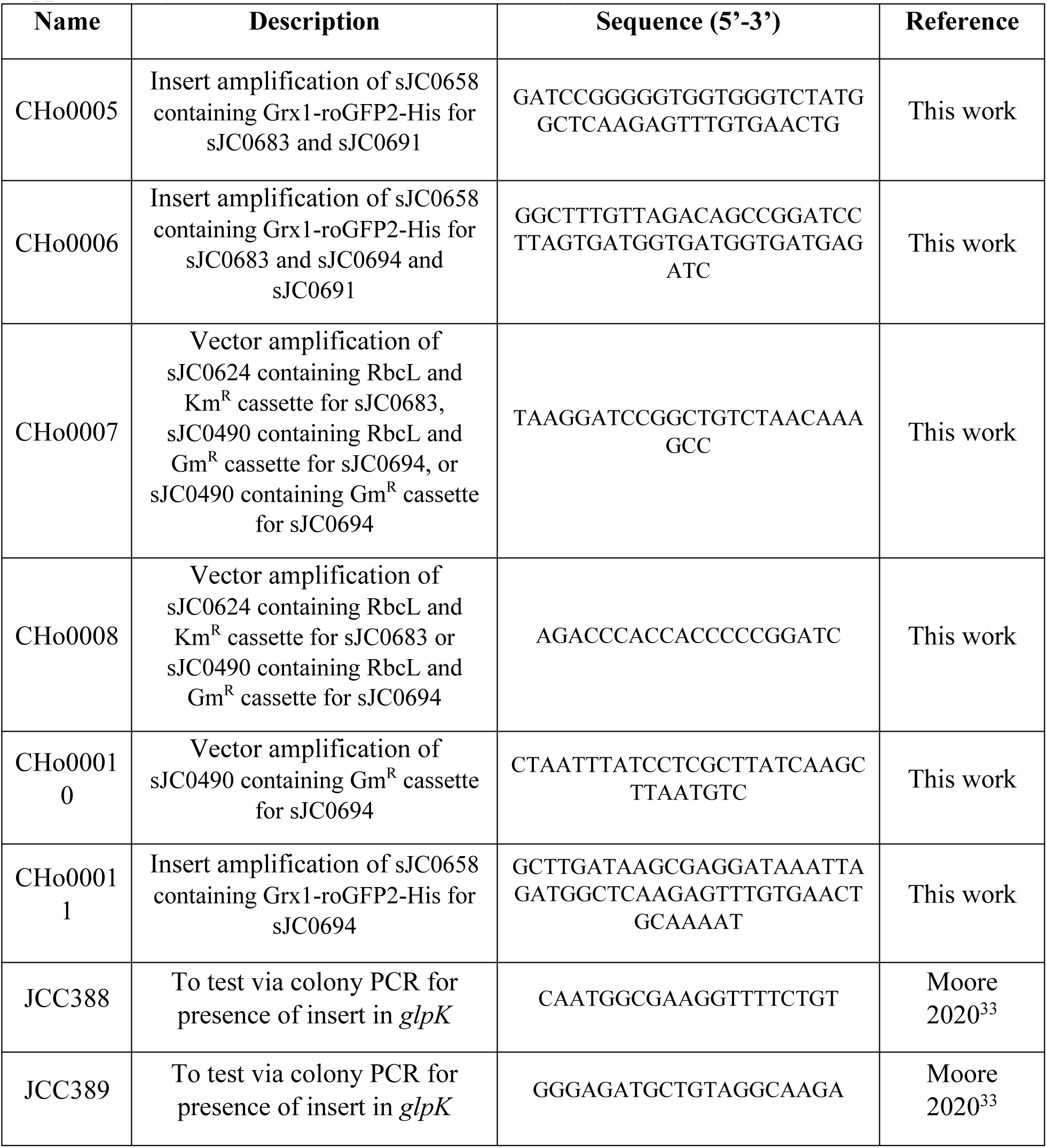
Primers used in this study.

**Supplementary Movie 1. Air Movie** Time-lapse imaging of roGFP expressing cells in brightfield, 395 nm (roGFP2), 640 nm (Cy5 for chlorophyll), 555 nm (RFP for phycobilisomes), and 470 nm (roGFP2). Cells were grown in air for the full duration of the movie.

**Supplementary Movie 2. Air Redox Movie** Time-lapse ratiometric images with *R*_395/470_ values ranging from 0 to 0.3. Cells were grown in air for the full duration of the movie. Similar results were observed in two additional independent experiments.

**Supplementary Movie 3. Air to 3% Movie** Time-lapse imaging of roGFP expressing cells in brightfield, 395 nm (roGFP2), 640 nm (Cy5 for chlorophyll), 555 nm (RFP for phycobilisomes), and 470 nm (roGFP2). Cells were grown in air for hours 0-12 and in 3% CO2 for hours 12-30.

**Supplementary Movie 4. Air to 3% CO_2_ Redox Movie** Time-lapse ratiometric images with *R*_395/470_ values ranging from 0 to 0.3. Cells were grown in air for hours 0-12 and in 3% CO2 for hours 12-30. Similar results were observed in two additional independent experiments.

**Supplementary Movie 5. 3% CO_2_ to Air Movie** Time-lapse imaging of roGFP expressing cells in brightfield, 395 nm (roGFP2), 640 nm (Cy5 for chlorophyll), 555 nm (RFP for phycobilisomes), and 470 nm (roGFP2). Cells were grown in 3% CO2 for hours 0-12 and in air for hours 12-30.

**Supplementary Movie 6. 3% CO_2_ to Air Redox Movie** Time-lapse ratiometric images with *R*_395/470_ values ranging from 0 to 0.3. Cells were grown in 3% CO2 for hours 0-12 and in air for hours 12-30. Similar results were observed in two additional independent experiments.

